# PrEditR: A protein-centric platform for CRISPR-mediated base editor sgRNA design

**DOI:** 10.64898/2026.05.15.725600

**Authors:** Felipe Vasquez-Castro, Leonardo D. Sanchez Solis, Samuel A. Myers

## Abstract

**Motivation:** Post-translational modifications (PTMs) are critical to protein activity, yet the biological role of most known modification sites remains uncharacterized. CRISPR-mediated phenotypic screens using base editors offer a powerful approach to dissecting PTM function at scale. However, existing sgRNA design tools for base editing applications are DNA-centric and lack the throughput required to integrate seamlessly with mass-spectrometry-based proteomics experimental outputs.

**Results:** We introduce protein editing in R, PrEditR, an open-source, protein-centric tool for highthroughput sgRNA design for custom base editor screens. PrEditR enables users to designate specific amino acid residues in proteins and design protospacer sequences that target the corresponding gene and annotate every resulting amino acid change (missense, nonsense, or silent), including bystander edits.

**Availability and Implementation:** PrEditR is available on GitHub and Docker Hub.

## Introduction

Post-translational modifications (PTMs) control nearly every aspect of cellular and organismal biology. Although mass spectrometry-based methods have mapped tens of thousands of chemically modified amino acid side chains across the proteome, our understanding of their roles and functions remain undetermined for more than 98% of over 220,000 sites reported in humans (Needham et al., 2019).

CRISPR-mediated base editors offer a promising genetic engineering tool to address this gap in knowledge. These tools expand the scope of genome editing by fusing a nickase Cas9 protein with specific nucleotide deaminase enzymes. Fusing Cas9 with a cytosine deaminase (e.g., rAPOBEC1) creates a Cytosine Base Editor (CBE), which drives cytosine-to-thymine (C-to-T) transitions. Conversely, fusion with an adenine deaminase (e.g., TadA) creates an Adenine Base Editor (ABE), resulting in adenine-to-guanine (A-to-G) transitions (Huang et al., 2021; Komor et al., 2016). Because base editors are built from Cas9 proteins, both CBEs and ABEs are programmable through single guide RNAs (sgRNAs) and are highly suitable for large-scale phenotypic screens. By designing a screen that specifically targets nucleotides encoding post-translationally modified amino acids, i.e. “knocking out” the PTM site without affecting transcriptional regulatory elements, base editors can be used to test the functional relevance of PTM sites in a high-throughput manner (Kennedy et al., 2024; Li et al., 2022).

Currently available tools to design sgRNAs for base editing are mostly DNA-centric, requiring users to provide a DNA sequence or genomic coordinates of the desired edit (Dinçer et al., 2026) instead of indicating a specific amino acid residue position in a given protein. Some are exclusively web-based, which limits their integration into automated pipelines (Chapdelaine-Trépanier et al., 2025; Hwang et al., 2018). Protein-level design has recently been addressed by tools such as BEscreen (Schneider et al., 2025), which allows targeting specific residues but requires users to specify the resulting mutation a priori (e.g., GAPDHL270F) and fails to generate a guide when the requested mutation is not feasible. Many proteins show a high degree of tolerance to random mutations, and in PTM functional screens, the identity of the substituted amino acid is often less important than the loss of the modifiable residue itself (Kennedy et al., 2024; Vila, 2025). To address these guide-design limitations, we developed PrEditR (pronounced “preditor”): an R-based tool for protein-centric sgRNA design for custom, reproducible base editing applications. PrEditR differs from prior protein-level tools in three respects that matter for PTM functional screens: it requires only the target residue, not the resulting mutation, and returns every sgRNA that edits it together with each guide’s resulting mutation(s); it is isoform-aware; and it is built for reproducible, offline, high-throughput batch use from local reference data, distributed as a multi-architecture Docker image with both a CLI and a GUI. The tool supports parallel execution and scales across computing environments, from personal laptops to high-performance clusters (HPCs), to meet the growing volume of data generated in proteomics experiments.

## Implementation

### Framework

PrEditR is written in R and containerized using Docker to ensure seamless cross-platform deployment. The Docker image is available for amd64 and arm64 architectures and offers both a command-line interface (CLI) and a Shiny-based graphical user interface (GUI) in the same image. To avoid performance bottlenecks from server-based requests, PrEditR reads genome assemblies, annotations, and identifier maps from local reference data rather than querying remote servers at run time. Reference data are distributed as modular, per-organism reference images to allow for adding support for additional organisms without modifying the application. The current registry provides eight organisms: *Homo sapiens* (GRCh38), *Mus musculus* (mm10), *Rattus novergicus* (rn7), *Danio rerio* (danRer11), *Drosophila melanogaster* (dm6), *Caenorhabditis elegans* (ce11), *Saccharomyces cerevisiae* (sacCer3), and *Gallus gallus* (galGal6). Optionally, users can provide a Bowtie-indexed genome, in .ebwt format, (Langmead et al., 2009) to perform protospacer off-target searches, reporting the number of genomic alignments at different mismatch levels. Parallel processing is supported via the future R framework (Bengtsson, 2021), allowing for computational scalability.

### Coordinate Mapping

PrEditR leverages the crisprVerse (Hoberecht et al., 2022) ecosystem and extends it with a bidirectional, splicing- and strand-aware translation between protein-residue space and genomic space (e.g., chr10:80786010-80786012 to valine 218 in transcript ENST00000264657), drawing genome sequences from UCSC/Bioconductor BSgenome packages, transcript models from Ensembl-derived annotation objects, and protein identifiers from local UniProt– Ensembl maps.

Each target is resolved to a specific Ensembl transcript using user-supplied Ensembl transcript ID or UniProt ID. Alternatively, PrEditR defaults to the canonical transcript when only a gene symbol is provided. Because every design is anchored to a specific transcript, PrEditR is isoform-aware: if the residue at the requested position on the mapped isoform does not match the requested amino acid, PrEditR halts that target and reports the inconsistency rather than generating a deceptive sgRNA.

Given the anchored transcript, a target residue is translated to its genomic codon coordinates, accounting for exon structure, strand orientation, and codons split across exon-intron junctions. This must be done in a perfectly precise manner because base editors typically have narrow optimal editing windows of around 4-10 bp (Qin et al., 2024). Candidate sgRNAs with a valid PAM are enumerated on either strand within the editing window, and the resulting edited sequence is mapped back in-frame to the protein to annotate every missense, nonsense, or silent change, including bystander edits. This back-translation, together with the split-codon and strand handling, is the step most error-prone in manual DNA-centric workflows as four combinations of gene-strand and guide-strand editing must be handled correctly: the target mutation can be generated directly on the coding strand or indirectly via the template strand, followed by corresponding endogenous repair on the coding strand for scenarios where the coding strand is located on either the positive or negative strand (Figure 1).

**Figure 1.**
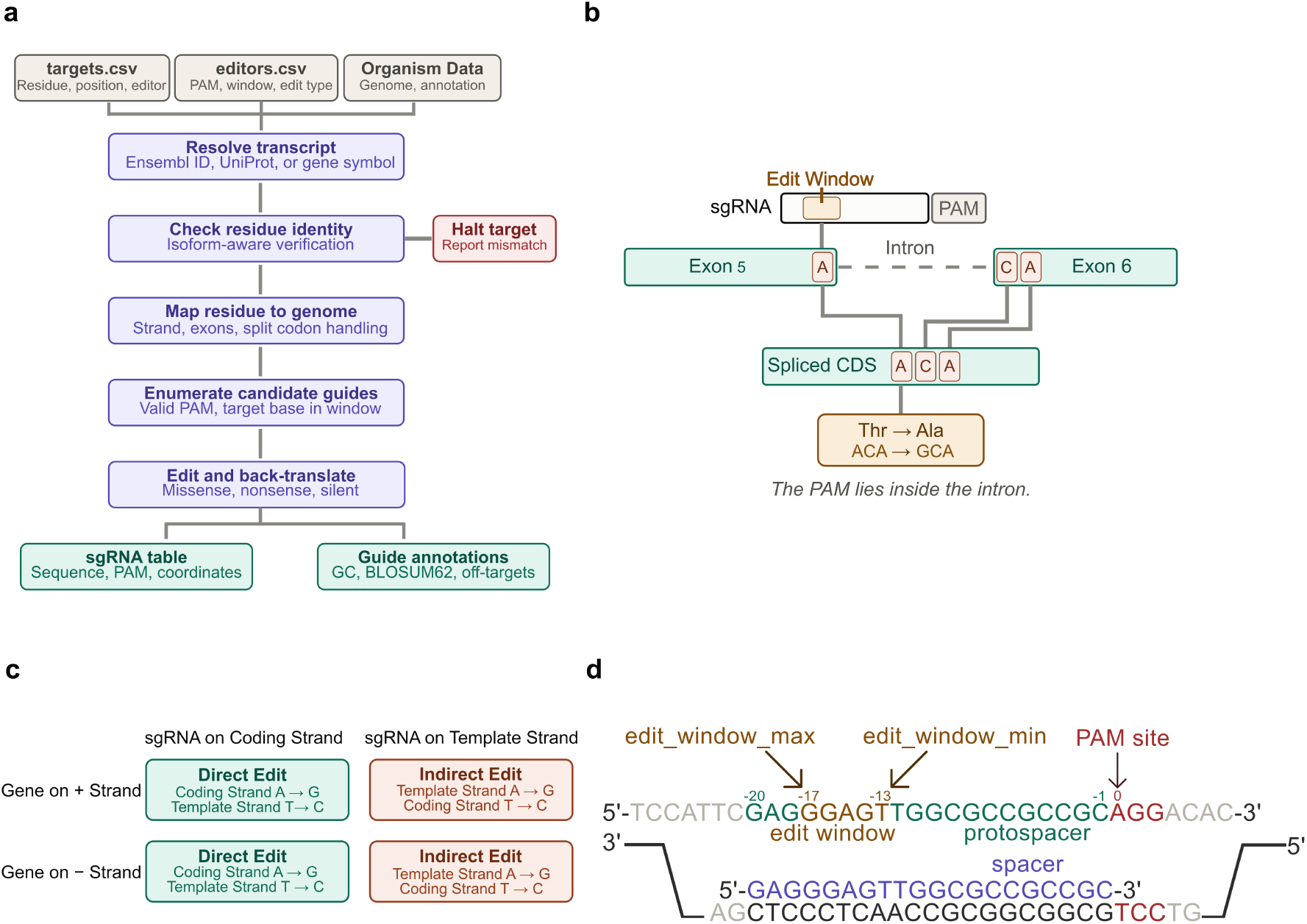
PrEditR design workflow and the coordinate logic it automates. (a) Design pipeline. Two configuration files (targets.csv, editors.csv) and a per-organism reference image supply all inputs. Each target is resolved to a single Ensembl transcript, supplied directly, mapped from a UniProt accession, or defaulted to the gene’s Ensembl canonical transcript. PrEditR verifies that the residue at the requested position on that isoform matches the requested amino acid; a mismatch halts the target and is flagged in the output rather than yielding a sgRNA. Verified residues are translated to genomic codon coordinates, accounting for exon structure, strand orientation, and codons split across exon–intron junctions. Candidate protospacers with a valid PAM are enumerated on either strand such that the target base falls within the editing window, and the edited sequence is back-translated in frame to annotate every missense, nonsense, and silent change, including bystander edits. sgRNA sequences, genomic coordinates for protospacer and PAM, and per-guide features (GC content, homopolymer and restriction-site flags, BLOSUM62 substitution score, and off-target alignment counts) are appended to the original input file.(b) A codon split across an exon–intron junction. The target adenine is the final base of exon 5; the remaining two bases of its codon lie at the start of exon 6. The protospacer and PAM therefore straddle the junction and sit partly within the intronic sequence, while the codon they alter is completed only after splicing. Here ACA (Thr) becomes GCA (Ala). Manual DNA-centric design requires two separate search windows to recover such guides; PrEditR resolves them in a single pass. (c) The four gene-strand × guide-strand combinations that determine the coding-strand outcome, shown for an adenine base editor. A protospacer matching the coding strand mutates the target adenine directly (A*→*G). A protospacer matching the template strand mutates the adenine paired with a coding-strand thymine, which endogenous repair resolves as T*→*C on the coding strand. For a cytosine base editor the indirect cells read G*→*A. (d) Editing window convention. Positions are numbered relative to the PAM, which is assigned position 0, with upstream (5*′*) positions taking negative values. edit_window_min is the position closest to the PAM and edit_window_max the position furthest from it. The window shown spans *−*13 to *−*17, the setting used for ABE8e throughout this work. Note that this convention differs from the protospacer numbering used in most published editor characterizations, which counts positions 1–20 from the PAM-distal end of the spacer. For a 20-nt sgRNA, standard position p corresponds to signed position *−*(21 *−* p), so a window reported as positions 4–8 is entered here as *−*17 to *−*13. This convention was selected to allow for sgRNAs of lengths different than 20 nt.

### Configuration

PrEditR’s pipeline can be run interactively (GUI) or from configuration files (GUI or CLI). The GUI’s “Search” tab lets users enter targets directly in a form and select editors from a built-in catalog. The editor picker is pre-populated with a curated catalog of validated base editors, each with its PAM, spacer length, conversion type, and editing window filled in and annotated with a literature and Addgene reference.

The “Batch Search” GUI tab and the CLI accept two configuration files for batch and automated workflows. The first of these files, the Base Editor Configuration (editors.csv), defines the properties of the base editors utilized for guide design and allows for multiple editors to be used in a single run. Each entry in this file requires a unique name as an identifier, the PAM sequence recognized by the nuclease (e.g., NGG, NG, NGN, …), and the protospacer length of the gRNA, which is typically 20 nucleotides. Additionally, this file must specify the specific nucleotide conversion (denoted as edit_type; e.g., “a2g” for an ABE) and the editing window through the *edit_window_min* and *edit_window_max values*, which represent the closest and furthest positions upstream of the PAM in nucleotides, respectively.

The second required file, the Targeting Specifications (targets.csv), details the amino acid targets and the specific base editors intended to modify them. A target may be identified by any one of a gene symbol, an Ensembl transcript ID, or a UniProt ID. A gene symbol alone resolves to the gene’s Ensembl canonical transcript. For each target, the file must specify the single-letter amino acid of the target residue and its numerical position within the protein sequence. Finally, the editor column must contain a name that exactly matches one of the unique identifiers defined in the editors.csv file to ensure the correct parameters are applied to the target site. Templates for these files are provided in Supplementary Tables 1 (targets.csv) and 2 (editors.csv), as well as the corresponding output example in Supplementary Table 3.

### Performance

To ensure PrEditR remains accessible to users without HPC resources, we optimized the platform to balance memory consumption and runtime across both small- and large-scale experiments. Parallel processing is driven by the future framework’s multicore strategy, which leverages process forking to efficiently share objects across workers. Because R’s native garbage collection might inadvertently disrupt this shared-memory model and inflate the memory footprint, we mitigate this overhead by aggressively minimizing the parent environment prior to parallel execution.

To analyze how peak memory scales with dataset size, we evaluated memory usage using the human phosphorylation site dataset from PhosphoSitePlus (phosphosite.org) (Hornbeck et al., 2012), splitting the data (240,184 phosphorylation sites) into 10% incremental steps from 10% to 100% at 24 threads. Peak proportional set size (PSS) grew from 34.8 GB at 10% of the dataset to 68.0 GB at 100% for a 10-fold increase in input size producing only a *∼*1.95-fold increase in memory. Thus, memory scales sublinearly over the range tested, PSS *≈* 30.4 GB + 0.147 MB/row (R^2^ = 0.96, n = 10 input sizes, 3 replicates each; Figure 2a). Thread scaling was then benchmarked on a fixed 10% subset of the same dataset (24,018 sites) using 4, 8, 12, 16, 24, 32, and 64 threads with 10 replicate runs per thread count, executed within a Singularity/Apptainer container on a system equipped with an AMD EPYC 7543P 32-core CPU (64 threads). As shown in Figure 2b, peak RAM usage scales linearly with the number of threads, rising from a mean PSS of 7.1 GB at 4 threads to 74.4 GB at 64 threads (*≈*1.1 GB per additional thread, R^2^ = 0.99), reflecting the *∼*1.2–1.5 GB working set held by each worker process. Mean wall-clock runtime fell from 11,852 s at 4 threads to 1,465 s at 64 threads to give an 8.1-fold speedup for a 16-fold increase in threads, with parallel efficiency declining from *∼*1.0 at 4–8 threads to 0.91 at 24 and 0.74 at 64, indicating diminishing returns once the allocation exceeds the 32 physical cores, consistent with a CPU-bound workload approaching hardware parallelism limits. To prevent double-counting shared objects, memory evaluation relied on peak proportional set size (PSS) rather than resident set size (RSS).

**Figure 2:**
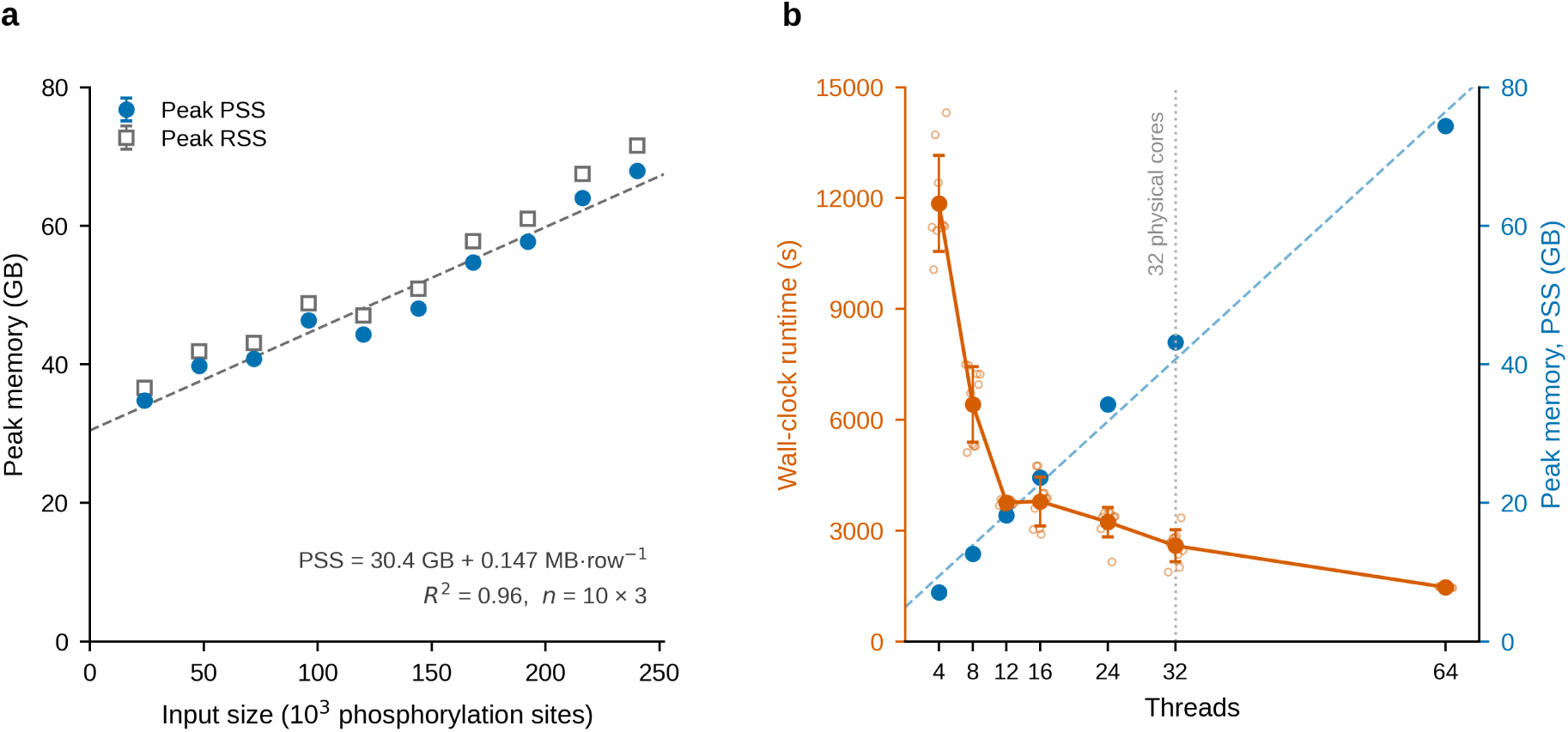
Memory and runtime scaling of PrEditR. All benchmarks were executed within a Singularity/Apptainer container on a single AMD EPYC 7543P node (32 physical cores, 64 hardware threads). Memory is reported as peak proportional set size (PSS) unless indicated otherwise, which avoids the double-counting of shared objects inherent to resident set size (RSS). (a) Peak memory as a function of input size. The human PhosphoSitePlus dataset (240,184 phosphorylation sites) was sampled in 10% increments from 10% to 100% and processed at a fixed 24 threads, with 3 replicate runs per input size (30 runs total). Blue filled circles, peak PSS; grey open squares, peak RSS; large markers with error bars denote mean *±* SD and small faint markers denote individual replicates. The dashed line is an ordinary least-squares fit to PSS across all runs (PSS = 30.4 GB + 0.147 MB·row*−*1; R^2^ = 0.96). (b) Wall-clock runtime (orange, left axis) and peak PSS (blue, right axis) as a function of thread count. A fixed 10% subset of the same dataset (24,018 sites) was processed using 4, 8, 12, 16, 24, 32, and 64 threads, with 10 replicate runs per thread count (70 runs total). Markers and error bars denote mean *±* SD; small open circles denote individual runtime replicates. The dashed blue line is a linear fit to peak PSS (PSS = 4.9 GB + 1.12 GB·thread*−*1; R^2^ = 0.99). The dotted vertical line marks the 32 physical cores of the test system. Runtime variability is greater at 4–32 threads (coefficient of variation 11–17%) than at 64 threads (*∼*1.4%), reflecting competition for resources from other jobs sharing the same compute nodes.

## Use Cases

### Functional Screening of Phosphorylation Sites

Kennedy et al. (2024) introduced a high-throughput approach to interrogate the functional relevance of phosphorylation sites and their control of NFAT:AP1 transcriptional activity by integrating base editing with phenotypic screening. Their target selection was informed by upstream phosphoproteomic data derived from mass spectrometry, which provides a protein-centric readout, i.e., UniProt accession number and modified amino acid number for a specific isoform. However, their sgRNA design followed a DNA-centric workflow. Specifically, the authors employed a custom script to generate all possible sgRNAs for every gene represented in their dataset. Subsequent annotation of the resulting amino acid mutations allowed them to filter for sgRNAs targeting phosphorylation sites identified in their phosphoproteomic analyses. This workflow resulted in the unnecessary generation of numerous guides, as it designed guides for entire genes despite only requiring a small subset relevant to the phosphosites of interest. Moreover, their pipeline was dependent on real-time queries to the Ensembl server, which may present limitations in scalability, particularly in large-scale screens where multiple base editors must be evaluated to identify the most effective one.

To evaluate the performance and robustness of PrEditR, we benchmarked the tool against the 9,200 A-to-G unique phosphorylation site targets from Kennedy et al. (2024) in Supplementary Table 2, which originally comprised 13,441 sgRNAs. Utilizing an AMD EPYC 7543P 32-core CPU with 24 threads, PrEditR successfully identified 11,451 sgRNAs (85.19%) by exact sequence identity (Supplementary Table 6). 1,978 sgRNAs (14.72%) were not found because the tool could not locate the target amino acid at the specified protein position. We cross-referenced the 1,978 missing sgRNAs with the original phosphoproteomics data from Supplementary Data 1 and confirmed that these discrepancies arose from mismappings between UniProt and Ensembl IDs. Specifically, for every failed guide represented in that subset, the Ensembl IDs reported in the sgRNA table did not correspond to the original UniProt IDs. PrEditR correctly identified a threonine at that position on the mapped isoform and halted the design upon finding an inconsistency between the desired target amino acid and the actual amino acid for that specific isoform. This type of inconsistency is noted in the PrEditR output.

### Functional Assessment of BRCA1 Variants

The base editor screen of BRCA1 variants by (Kweon et al., 2020) serves as a good use case to demonstrate the significant workflow improvements offered by PrEditR when creating an amino acid-targeting mutational screen. The original method required manual retrieval of the large BRCA1 exon and intron-exon junction sequences. The 23 exons and their corresponding junctions with introns were fragmented into segments under 1000 bp for submission to CasDesigner (Park et al., 2015). PrEditR replaces this multi-step, laborious process with a single run to design guides targeting all amino acids in the protein.

We evaluated PrEditR against the 1,863 residues of BRCA1 (UniProt P38398-1; Ensembl ENST00000357654). Using 30 threads and a C-to-T editor, PrEditR identified 465 guides targeting 360 residues (Supplementary Table 5) in 225 seconds. While the original study did not provide the full output of their sgRNA search across the entire BRCA1 gene, comparison with their screened guides on Supplementary Table 1 showed that PrEditR recapitulated all (100%) sgRNA sequences. Furthermore, PrEditR correctly reported the global translational effects of multiple base edits within single codons. For instance, in exon 23, where the original authors annotated individual G1803D (c.5408G>A) and G1803S (c.5407G>A) mutations, PrEditR accurately consolidated these adjacent edits into the resulting G1803N substitution.

## Validation

### Stratified Validation Set Generation

To evaluate PrEditR’s accuracy and scalability, we designed ABE-NGG guides for 1,003 target protein-encoding genes using a base editing window of −13 to −17, resulting in an initial dataset of 974,003 target evaluations. This dataset encompassed both residues with successfully designed sgRNAs, and residues predicted to be untargetable. To construct a validation set that thoroughly tested PrEditR across diverse genomic contexts, we implemented a stratified sampling approach. We selected 72 unique target residues, taking one representative residue for every possible combination of four parameters: guide availability (none, one, or multiple guides per residue), transcript region (first, second, or final third of the gene coding sequence), target strand orientation (coding or template), and exon boundary proximity (exon-internal, exon start edge, exon end edge, or split across an intron-exon junction). As such, we established a final validation set of 106 test cases, comprising 82 predicted sgRNAs and 24 instances where the tool identified a residue as untargetable, providing a comprehensive suite of edge cases to guide the refinement of the tool’s underlying algorithms.

### Manual Benchmarking and Concordance

We performed manual sgRNA design for each target in the validation set using commonly utilized DNA-centric web tools. For each protein of interest, its relevant transcript was obtained from Ensembl and imported into Benchling. We utilized a 45-bp search window centered on the target residue to identify valid sgRNAs via the RGEN BE-Designer tool (Hwang et al., 2018). For codons split across exons, we utilized dual search windows centered around each segment of the split codon to ensure the manual verification captured the full range of potential residue targeting. In addition, all editable bases within the base editing window were assumed to be edited.

This manual-guided debugging allowed us to resolve logic errors in complex genomic scenarios, ultimately achieving 100% concordance. PrEditR successfully identified all 82 valid sgRNAs, correctly predicted all 24 untargetable instances, and accurately annotated every resulting mutation across the entire validation set.

## Discussion

Base editors have revolutionized the molecular biology toolbox by enabling targeted nucleotide conversions within a narrow window, allowing researchers to investigate PTM function at an unprecedented scale. However, a significant gap remains for reproducible, largescale sgRNA design. PrEditR fulfills this need as a “one-stop” solution that streamlines sgRNA design while integrating critical experimental and analytical parameters often neglected by existing software.

PrEditR is a design tool for both hypothesis-driven and library-scale PTM screens on the same code path; per-target filtering (guide availability, mutation type, built-in controls) itself aids library-size management, while feasibility constraints such as synthesis limits, delivery, and readout remain the user’s experimental choices. Gene-symbol input and the form-based GUI make routine design accessible to experimental biologists and proteomics researchers, while the CLI and reference-image system serve bioinformaticians building large or automated libraries.

Beyond simple design, the tool enhances experimental reliability by screening for downstream restriction enzyme sites and incorporating essential non-targeting and non-editing controls that are critical for establishing baselines for screens. For guide prioritization, PrEditR reports editor-agnostic sequence features such as GC content, homopolymer/repeat flags, restriction-site presence flags, and a BLOSUM62 substitution score (Henikoff and Henikoff, 1992) per predicted amino acid change. When a Bowtie-indexed genome is supplied, PrEditR reports the genomic alignment counts at various mismatch levels. An optional –dna_context flag appends the editing-window sequence plus 50 nt of upstream and downstream flanking sequence (dna_context_upstream, dna_edit_window, dna_context_downstream), so users can feed sequence context to an editor-matched efficiency-prediction model of their choice. We deliberately do not embed such a model because published models are editor-, context-, and cell-type-specific and validated for only a few editors. PrEditR also provides exact genomic start and end coordinates for both the protospacer and the PAM of every guide, which facilitates manual verification when necessary. Efficiency is further improved through an “append-to-input” output structure; by preserving and attaching results to the original input file, the tool allows researchers to include their own metadata that remains intact for downstream use.

The editing window is a per-editor parameter, so PrEditR represents both the narrow windows of early ABEs (ABE7.10, ABEmax) and the expanded windows of high-activity variants (ABE8e, ABE8e-V106W) simply by configuring the appropriate bounds. The curated editor catalog already includes ABE8e, and our validation used a *−*13 to *−*17 window. PrEditR does not model position-specific editing probabilities, so for broad windows bystander-edit predictions are conservative.

PrEditR’s recapitulation of the independently designed and experimentally recovered BRCA1 (Kweon et al., 2020) and phosphosite-targeting (Kennedy et al., 2024) guides, together with the 100% concordance on our stratified manually verified validation set, provides indirect experimental support for its designs.

To maximize accessibility, we developed PrEditR with both a command-line interface and an intuitive GUI. The software underwent extensive RAM optimization, democratizing access to researchers that lack high-performance workstations. Finally, by providing Docker images for both arm64 and amd64 architectures, we ensure that PrEditR is portable and accessible across the full spectrum of modern computing hardware.

## Limitations and Troubleshooting

1. PrEditR assumes that each base editor performs a single type of nucleotide mutation (e.g., A-to-G, C-to-T, C-to-G, …). For editors capable of multiple conversion types, a workaround is to define them as separate editor entries, one for each distinct conversion.
2. Only DNA base editors are supported.
3. PAM sequences are assumed to lie immediately downstream (i.e., 3’) of the protospacer.
4. All protospacers designed in a single run must be of the same length.
5. PrEditR assumes uniform editing efficiency across all positions within the defined editing window; position-specific editing probabilities (i.e., weighted editing windows) are currently not supported.
6. PrEditR reads reference data (genome, annotation, and identifier maps) from per-organism reference images. The current registry provides *Homo sapiens*, *Mus musculus*, *Rattus norvegicus*, *Danio rerio*, *Drosophila melanogaster*, *Caenorhabditis elegans*, *Saccharomyces cerevisiae*, and *Gallus gallus*. To prevent server-querying-related bottlenecks, local UniProt-to-Ensembl ID maps were constructed from Ensembl as of November 25, 2025. These mappings might change over time, especially as unreviewed UniProt entries are updated. Because reference data are modular, additional organisms or updated maps can be supplied without modifying the application.
7. PrEditR queries local genetic databases using Ensembl IDs. If only a gene symbol or UniProt ID is provided, PrEditR will first map the supplied ID to the corresponding Ensembl ID using the built-in database. For novel UniProt to Ensembl transcript ID associations (i.e., those identified after November 25, 2025), the local database might fail to find the corresponding Ensembl transcript ID provided only a gene symbol or UniProt ID. Providing the transcript Ensembl ID instead will circumvent this error.
8. The graphical interface offers a curated catalog of validated editors. User-defined editors are supported for custom or novel applications but are only as reliable as the parameters provided, and outcomes for editors without published characterization should be interpreted with caution.

## Acknowledgements

We thank members of the Myers lab and Dr. Patrick Hogan for their valuable input. We are grateful to Dr. Terry Gaasterland for her guidance.

## Supplementary data

Supplementary data accompany this manuscript.

## Data availability

The source code and full documentation for PrEditR are available on GitHub (https://github.com/fvasquezcastro/preditr). PrEditR is distributed via Docker Hub (https://hub.docker.com/repository/docker/fvasquezcastro/preditr/general).

## Author contributions

FVC: Conceptualization (Supporting), Methodology (Lead), Software, Writing – original draft (Lead), Writing – review & editing (Lead). LSS: Methodology (Supporting), Validation, Writing – original draft (Supporting), Writing – review & editing (Supporting). SAM: Conceptualization (Lead), Methodology (Supporting), Writing – review & editing (Equal), Funding acquisition.

## References

Bengtsson, H. (2021). A Unifying Framework for Parallel and Distributed Processing in R using Futures. The R Journal, 13(2), 208. 10.32614/RJ-2021-048

Chapdelaine-Trépanier, V., Shenoy, S., Masud, W., Minju-OP, A., Bérubé, M.-A., Schönherr, S., Forer, L., Fradet-Turcotte, A., Taliun, D., & Cuella-Martin, R. (2025). CRISPR-BEasy: a free web-based service for designing sgRNA tiling libraries for CRISPR-dependent base editing screens. Nucleic Acids Research, 53(W1), W193–W202. 10.1093/nar/gkaf382

Dinçer, C., Fussing, B., Garnett, M. J., & Coelho, M. A. (2026). BEstimate: a computational tool for the design and interpretation of CRISPR base editing experiments. Genome Biology, 27(1). 10.1186/s13059-026-04077-z

Henikoff, S., & Henikoff, J. G. (1992). Amino acid substitution matrices from protein blocks. Proceedings of the National Academy of Sciences of the United States of America, 89(22), 10915–10919. 10.1073/pnas.89.22.10915

Hoberecht, L., Perampalam, P., Lun, A., & Fortin, J.-P. (2022). A comprehensive Bioconductor ecosystem for the design of CRISPR guide RNAs across nucleases and technologies. Nature Communications, 13(1), 6568. 10.1038/s41467-022-34320-7

Hornbeck, P. V., Kornhauser, J. M., Tkachev, S., Zhang, B., Skrzypek, E., Murray, B., Latham, V., & Sullivan, M. (2012). PhosphoSitePlus: a comprehensive resource for investigating the structure and function of experimentally determined post-translational modifications in man and mouse. Nucleic Acids Research, 40(Database issue), D261–D270. 10.1093/nar/gkr1122

Huang, T. P., Newby, G. A., & Liu, D. R. (2021). Precision genome editing using cytosine and adenine base editors in mammalian cells. Nature Protocols, 16(2), 1089–1128. 10.1038/s41596-020-00450-9

Hwang, G.-H., Park, J., Lim, K., Kim, S., Yu, J., Yu, E., Kim, S.-T., Eils, R., Kim, J.-S., & Bae, S. (2018). Web-based design and analysis tools for CRISPR base editing. BMC Bioinformatics, 19(1), 542. 10.1186/s12859-018-2585-4

Kennedy, P. H., Alborzian Deh Sheikh, A., Balakar, M., Jones, A. C., Olive, M. E., Hegde, M., Matias, M. I., Pirete, N., Burt, R., Levy, J., Little, T., Hogan, P. G., Liu, D. R., Doench, J. G., Newton, A. C., Gottschalk, R. A., De Boer, C. G., Alarcón, S., Newby, G. A., & Myers, S. A. (2024). Post-translational modification-centric base editor screens to assess phosphorylation site functionality in high throughput. Nature Methods, 21(6), 1033–1043. 10.1038/s41592-024-02256-z

Komor, A. C., Kim, Y. B., Packer, M. S., Zuris, J. A., & Liu, D. R. (2016). Programmable editing of a target base in genomic DNA without double-stranded DNA cleavage. Nature, 533(7603), 420–424. 10.1038/nature17946

Kweon, J., Jang, A.-H., Shin, H. R., See, J.-E., Lee, W., Lee, J. W., Chang, S., Kim, K., & Kim, Y. (2020). A CRISPR-based base-editing screen for the functional assessment of BRCA1 variants. Oncogene, 39(1), 30–35. 10.1038/s41388-019-0968-2

Langmead, B., Trapnell, C., Pop, M., & Salzberg, S. L. (2009). Ultrafast and memory-efficient alignment of short DNA sequences to the human genome. Genome Biology, 10(3), R25. 10.1186/gb-2009-10-3-r25

Li, J., Lin, J., Huang, S., Li, M., Yu, W., Zhao, Y., Guo, J., Zhang, P., Huang, X., & Qiao, Y. (2022). Functional Phosphoproteomics in Cancer Chemoresistance Using CRISPR-Mediated Base Editors. Advanced Science (Weinheim, Baden-Wurttemberg, Germany), 9(30), 2200717. 10.1002/advs.202200717

Needham, E. J., Parker, B. L., Burykin, T., James, D. E., & Humphrey, S. J. (2019). Illuminating the dark phosphoproteome. Science Signaling, 12(565), eaau8645. 10.1126/scisignal.aau8645

Park, J., Bae, S., & Kim, J.-S. (2015). Cas-Designer: a web-based tool for choice of CRISPRCas9 target sites. Bioinformatics (Oxford, England), 31(24), 4014–4016. 10.1093/bioinformatics/btv537

Qin, W., Liang, F., Lin, S.-J., Petree, C., Huang, K., Zhang, Y., Li, L., Varshney, P., Mourrain, P., Liu, Y., & Varshney, G. K. (2024). ABE-ultramax for high-efficiency biallelic adenine base editing in zebrafish. Nature Communications, 15(1). 10.1038/s41467-024-49943-1

Schneider, P. G., Liu, S., Bullinger, L., & Ostendorf, B. N. (2025). BEscreen: a versatile toolkit to design base editing libraries. Nucleic Acids Research, 53(W1), W68–W72. 10.1093/nar/gkaf406

Vila, J. A. (2025). Physical principles underpinning molecular-level protein evolution. European Biophysics Journal, 54(5), 201–211. 10.1007/s00249-025-01776-6

